# Mechanistic analysis of the SERK3 *elongated* allele defines a role for BIR ectodomains in brassinosteroid signaling

**DOI:** 10.1101/257543

**Authors:** Ulrich Hohmann, Joël Nicolet, Andrea Moretti, Ludwig A. Hothorn, Michael Hothorn

## Abstract

The leucine-rich repeat receptor kinase (LRR-RK) BRI1 requires a shape-complementary SERK co-receptor for brassinosteroid sensing and receptor activation. Interface mutations that weaken the interaction between receptor and co-receptor *in vitro* reduce brassinosteroid signaling responses. The SERK3 elongated (*elg*) allele maps to the complex interface and shows enhanced brassinosteroid signaling, but surprisingly no tighter binding to the BRI1 ectodomain *in vitro.* Here, we report that rather than promoting the interaction with BRI1, the *elg* mutation disrupts the ability of the co-receptor to interact with the ectodomains of BIR receptor pseudokinases, negative regulators of LRR-RK signaling. A conserved lateral surface patch in BIR LRR domains is required for targeting SERK co-receptors and the *elg* allele maps to the core of the complex interface in a 1.25 Å BIR3 - SERK1 structure. Collectively, our structural, quantitative biochemical and genetic analyses suggest that brassinosteroid signaling complex formation is negatively regulated by BIR receptor ectodomains.

## Main

The LRR-RK BRASSINOSTEROID INSENSITIVE 1 (BRI1) is the major receptor for growth-promoting steroid hormones in plants^1,2^ and binds brassinosteroids (BRs) including the potent brassinolide (BL) with its LRR ectodomain^3,4^. Ligand-associated BRI1 can interact with the LRR domain of a SOMATIC EMBRYOGENESIS RECEPTOR KINASE (SERK) co-receptor kinase, which completes the steroid binding site^5,6^. Heterodimerisation of the receptor and co-receptor LRR domains at the cell surface enables the kinase domains of BRI1 and SERK to trans-phosphorylate each other, allowing BRI1 to activate the cytoplasmic side of the brassinosteroid signaling cascade^7-9^. Mutations in the BRI1 - SERK complex interface that reduce binding between the receptor and co-receptor ectodomains *in vitro*, weaken the interactions of the full-length proteins *in planta* and consequently result in BR loss-of-function phenotypes^10^. Previously, two gain-of-function mutations have been reported for the BR signaling complex. The BRI1 *sud1* allele stabilizes the steroid binding site of the receptor^11,5^. A similar phenotype is observed with the *elg* mutant^12^, originally identified as a suppressor of *ga4*, a gibberellic acid biosynthetic enzyme^13^. SERK3^D122^ is replaced by an asparagine residue in *elg* mutant plants^14^ and Asn122 maps to the constitutive BRI1 - SERK3 complex interface outside the steroid binding pocket^5,6,10^ (Fig. 1a). In BRI1 - SERK complex structures, SERK3^D122^ stabilizes the conformation of SERK3^R146^, which in turn makes polar contacts with BRI1^E749 3,5,10^ (Fig. 1a). Mutation of the corresponding Asp128 to asparagine in rice SERK2 has been shown to alter these interactions^15^. SERK3^D122^ also positions SERK3^E98^ to allow for interaction with BRI1^T750^, which is found replaced by isoleucine in *bri1-102* loss-of-function mutants^16^ (Fig. 1a). Taken together, SERK3^D122^ is in contact with several residues critically involved in BR signaling complex formation.

**Fig. 1:**
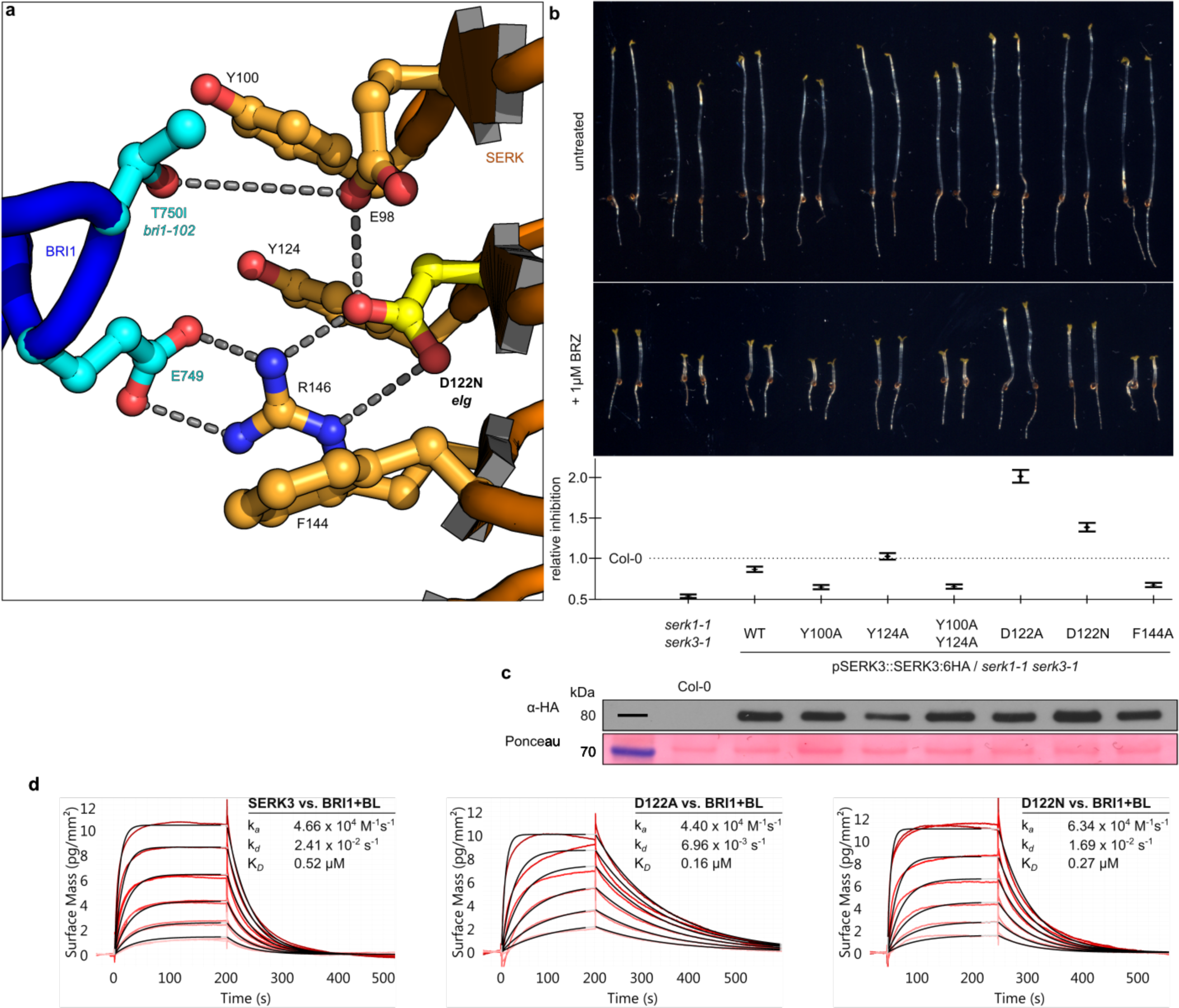
SERK3 elg is a gain of function allele *in vivo* but not *in vitro.* **a**, Ribbon diagram of the *elg*-containing complex interface, as seen in the BRI1 – BL – SERK1 structure (PDB-ID 4LSX^5^). BRI1 and SERK are depicted in blue and orange, respectively, selected residues are shown in ball-and-stick representation with the elg residue Asp122 highlighted in yellow. Polar interactions are shown as dotted lines. SERK residue numbering is according to the SERK3 sequence throughout. **b**, Hypocotyl growth assay of dark grown seedlings in the pre- and absence of the BR biosynthesis inhibitor brassinazole (BRZ). The BRZ hypersensitivity seen in the *serk1-1 serk3-1* mutant is complemented by the expression of SERK3^WT^ (Col-0 is the untransformed wild-type). Shown alongside is the quantification of the data with relative inhibition plotted together with lower and upper confidence intervals (N=5, n=50). **c**, Western blot using an HA antibody against SERK3:HA from plant material shown in (**a**). A Ponceau loading control is shown alongside. **d**, Binding kinetics for SERK3, SERK3^D122A^ and SERK3^D122N^ (*elg*) vs. BRI1 in the presence of BL obtained from grating-coupled interferometry (GCI) experiments. Sensograms with recorded data are shown in red with the respective fits in black, and include table summaries of the corresponding association rate constant (k_a_), dissociation rate constant (k_*d*_) and dissociation constant K_*D*_.

We complemented a *serk1-1 serk3-1* double mutant with 6xHA-tagged wild-type or SERK3 mutant genomic constructs under the control of the SERK3 promoter. We could recapitulate the gain-of-function phenotype of SERK3^D122N^ plants in quantitative hypocotyl growth assays^12^ and replacing SERK3^D122^ with alanine resulted in an even stronger BR signaling phenotype (Figs. 1b,c S1, TableS1). We produced SERK3^D122N^ and SERK3^D122A^ LRR domains by secreted expression in insect cells and characterized their interaction with the BRI1 ectodomain in grating-coupled interferometry (GCI) binding assays^10^. The binding kinetics reveal that wild-type and mutant SERK3 LRR domains bind BRI1 with similar association rates (k_a_) (Fig. 1d). SERK3^D122A^ but not SERK3^D122N^ has a slower dissociation rate (kd) from the receptor, and consequently a slightly lower dissociation constant (K_D_). Overall, the only moderately altered binding kinetics for wild-type vs. mutant SERK3 ectodomains cannot rationalize their gain-of-function phenotype *in planta* (Fig. 1b-d).

Recently, the BRI1-ASSOCIATED-KINASE1 INTERACTING KINASE 3 (BIR3) has been reported as a negative regulator of BR signaling in Arabidopsis^17^. Ectopic overexpression of BIR3 results in BR loss-of-function phenotypes including BL insensitivity and reduced BRI1-EMS-SUPPRESSOR 1 (BES1) dephosphorylation^17^. The cytosolic pseudokinase domains of BIR2 and BIR3 bind the SERK3 kinase domain in yeast-2-hybrid assays and the full-length proteins interact *in planta*^17,18^. We hypothesized that also the highly conserved BIR ectodomains may contribute to BIR3 - SERK3 complex formation. Indeed, we found that the recombinantly purified BIR3 LRR domain binds SERK3 with a K_D_ of ~1 μM and with 1:1 stoichiometry (N) in isothermal titration calorimetry (ITC) experiments (Fig. 2a). No binding was detected between the BIR3 and BRI1 ectodomains (Fig. 2a). While BIR3 and BIR2 cannot discriminate between different SERK ectodomains *in vitro* (K_D_ ranges from ~1 to ~3 μM), *bir3* but not *bir2-1* or *bir2-3* mutant plants display a weak BR gain-of-function signaling phenotype (Figs. 2a-c, S1,S3, Table S1). SERK - BIR complex formation is likely driven by their extracellular LRR domains, as we could not observe detectable binding of the cytoplasmic (pseudo)kinase domains in ITC assays (Fig. S2).

**Fig. 2:**
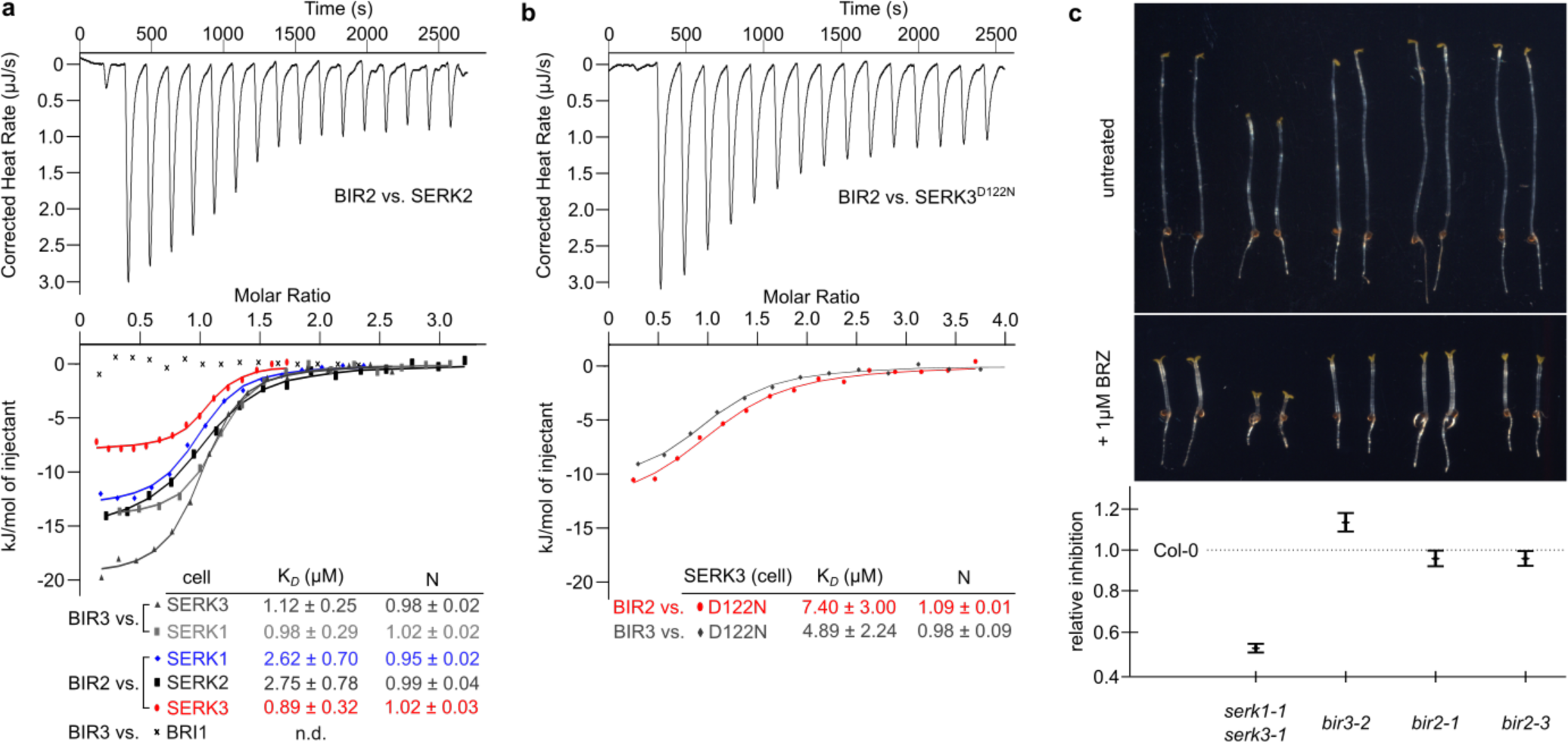
BIR ectodomains interact with different SERK co-receptors *in vitro.* **a,b**, Isothermal titration calorimetry (ITC) experiments of BIR2 and BIR3 LRR domains vs. (**a**) wild-type SERK ectodomains and (**b**) vs. the SERK3^D122N^ mutant ectodomain and including table summaries for dissociation constants (K_D_,) and binding stoichiometries (N) (± fitting error; n.d.: no detectable binding). **c**, Hypocotyl growth assay in the pre- and absence of BRZ (compare Fig. 1b). Relative inhibition together with upper and lower confidence intervals are shown alongside; Col-0 and *serk1-1 serk3-1* are the same as shown in Fig. 1b (N=5, n=50).

We next tested, if the *elg* mutation could modulate the interaction between BIRs and SERK3. Indeed, the SERK3^D122N^ mutant shows ~4-fold reduced binding to BIR3 and ~8-fold reduced binding to BIR2 (Fig. 2a,b). Due to its low expression yield, the SERK3^D122A^ mutant (Fig. 1) could not be assayed by ITC. Together, our experiments suggest that SERK3^D122^ maps to the interface of different SERK3 - BIR complexes and that interactions between interface residues may be compromised in the *elg* mutant background.

To gain insight into the BIR targeting mechanism, we sought to determine a crystal structure of BIR3 but did not succeed in obtaining diffraction quality crystals. Crystals of the related BIR2 ectodomain (residues 29-221, ~60% sequence identity with BIR3) diffracted to 1.9 Å resolution (Table S2). BIR2 contains five LRRs and shows a high degree of structural conservation with SERKs (r.m.s.d is ~1.5 Å comparing 175 corresponding C_a_ atoms in BIR2 and SERK1) with the exception of a protruding loop in the N-terminal capping domain of BIR2 (magenta in Fig. 3a). The BIR2 N- and C-terminal caps as well as the LRR core are stabilized by disulfide bridges conserved among the different BIR family members (Figs. 3c, S4). The conserved Asn58 in the BIR2 N-cap is glycosylated in our structure (Fig. 3c, S4). A set of solvent exposed hydrophobic residues including BIR2^W73^ from the protruding loop, BIR2^F128^, BIR2^F152^ and BIR2^R176^ form a lateral surface patch conserved among BIRs from different species, but not in SERK proteins (Figs. 3b,c, S4). This potential interaction surface differs from the central binding platform used by SERKs for targeting ligand-sensing LRR-RKs (Fig. 3c)^9,10^. We generated several point-mutations in the respective surface areas and assayed the mutant proteins vs. SERK3 in ITC assays. BIR2^E84R^ and BIR2^V157D^ originating from the central LRR groove still bind SERK3, suggesting that this interaction platform is not used by BIRs to target SERKs (Figs. 3c,d). Mutation of BIR2^W73^ from the protruding N-cap loop to alanine weakens the interaction with SERK3 and replacing BIR2^F152^ or BIR2^R176^ from the lateral surface patch with alanine disrupts binding (Figs. 3c,d). Thus, the unique N-cap loop and the lateral surface patch in the LRR domain of BIR2 are involved in the interaction with SERK3.

**Fig. 3:**
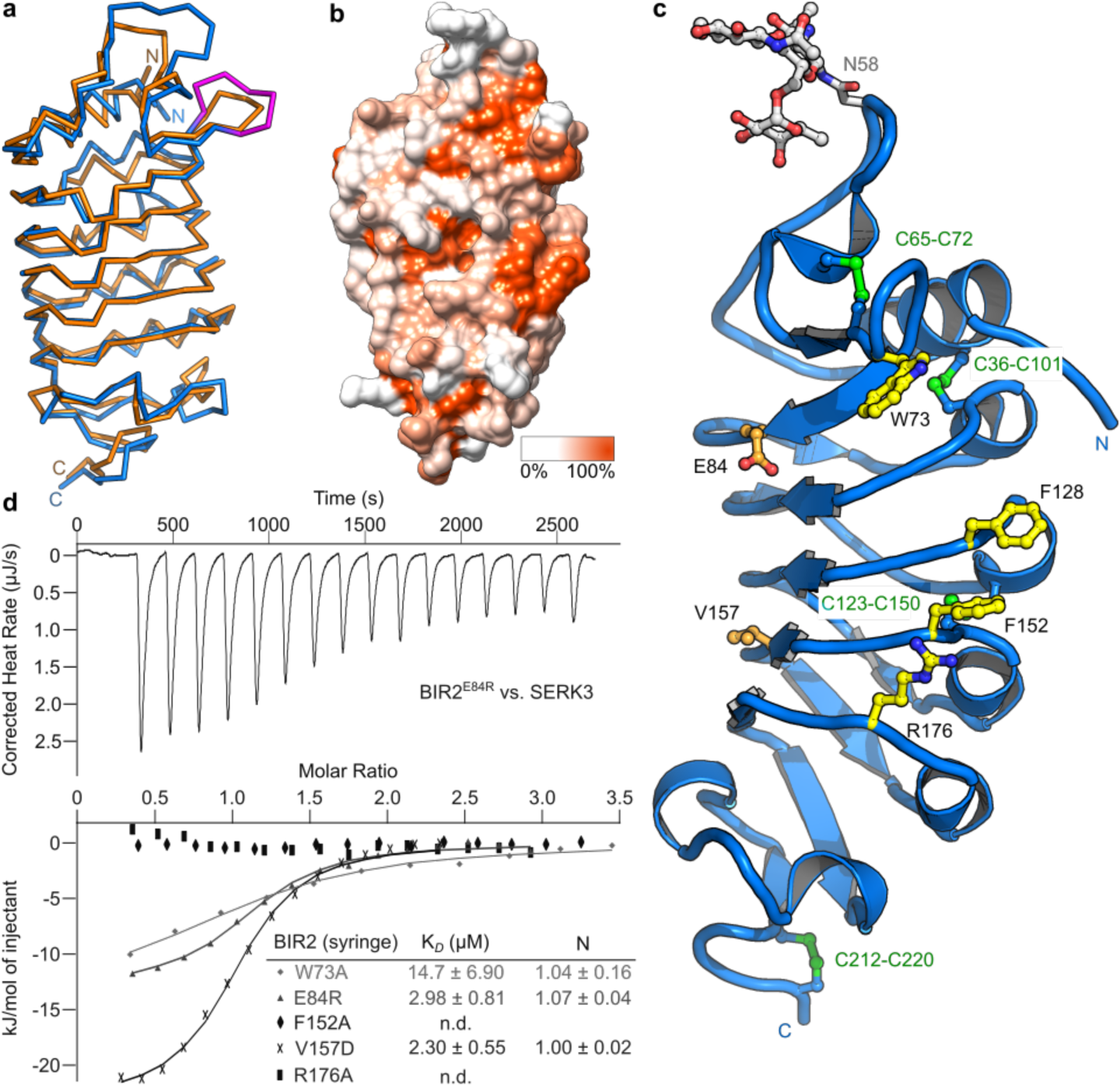
The BIR2 ectodomain adopts a SERK-like fold with an additional lateral protein interaction interface. **a**, Structural superposition of the isolated BIR2 and SERK1 (PDB-ID 4LSC^5^) ectodomains (r.m.s.d. is ~1.5 Å comparing 175 corresponding C_α_ atoms). C_α_ traces of SERK1 (orange) and BIR2 (blue) are shown; the unique, protruding BIR2 N-terminal cap loop region is highlighted in magenta. **b**, Surface representation of the BIR2 ectodomain, gradient colored according to the amino-acid sequence conservation of BIR proteins from different species (compare Fig. S4). **c**, The extracellular BIR2 domain consists of five LRRs with N- and C-terminal capping domains and a lateral protein interaction interface. Shown is a ribbon diagram of the BIR2 LRR domain (in blue), the four disulfide bonds are highlighted in green, selected residues in the lateral interface are in yellow, residues in the LRR central groove in orange, and the N-glycan moiety in gray (all in ball-and-sticks representation). **d**, ITC experiments of BIR2 ectodomain mutants vs. the extracellular domain of SERK3 with table summaries alongside.

To understand how BIRs target the central, *elg*-containing surface in SERKs, we performed crystallization trials for various BIR - SERK ectodomain combinations. We obtained crystals for a BIR3 - SERK1 complex diffracting to 1.25 Å resolution (Table S2). Our crystals contain a fully glycosylated BIR3 - SERK1 heterodimer in the asymmetric unit, consistent with their in solution behavior (Figs. 4a, S5). Most surface areas of the SERK1 LRR domain are shielded by carbohydrate, except for the central interaction surface used to, for example, bind the BRI1 and HAESA ligand-sensing LRR-RKs^3,5,10,19^. Structural comparison of the BIR3 - SERK1 complex with structures of the isolated SERK1 and BIR2 ectodomains reveals no major conformational rearrangements in BIRs and SERKs upon complex formation, with the exception of the protruding loop containing BIR2^W73^ or the corresponding Trp67 in BIR3 (Fig. S6). In the complex structure, BIR3 establishes a network of hydrophobic and polar interactions with the SERK1 C-terminal cap and with the two C-terminal LRRs (total buried complex surface area is ~1,400 Å^2^) (Fig. 4a). Several polar contacts are mediated by water molecules. The complex structure reveals the tip of the BIR3 protruding N-cap loop in direct contact with the SERK1 *elg* surface (Fig. 4b). SERK residues Asp122 (numbering corresponds to SERK3 throughout) and the neighboring Tyr124 together coordinate a water molecule, which in turn hydrogen bonds with BIR3^E69^ in the protruding loop tip (Fig. 4b). The neighboring Tyr100 establishes an additional hydrogen bond with BIR3^E69^ and the remaining loop tip residues BIR3^N68^ and BIR3^K70^ form similar interaction with SERK residues Asn148 and Asn77, respectively (Fig. 4b). Importantly, mutation of SERK Tyr100 or Tyr124 to alanine reduces BIR2 binding (Fig. 4b,d). Both tyrosine residues are also part of the BRI1 - SERK complex interface and, importantly, mutation of SERK3^Y100^ but not SERK3^Y124^ to alanine weakens the interaction with BRI1 (Fig. 4e).

**Fig. 4:**
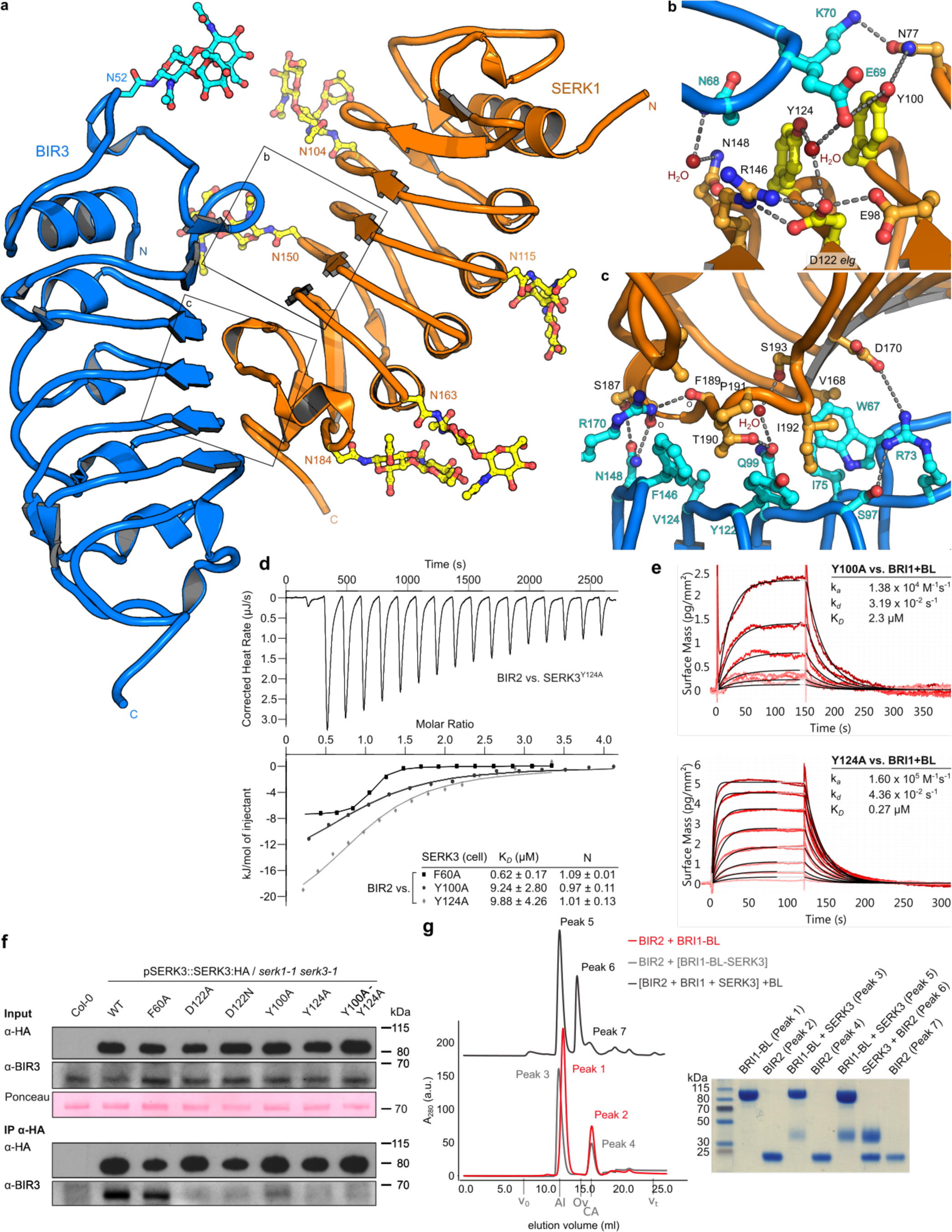
A BIR3-SERK1 complex structure provides a mechanism for SERK gain-of-function mutations. **a**, Structure of the BIR3 – SERK1 ectodomain complex, with BIR3 shown in blue and SERK1 in orange and with N-glycans highlighted in ball-and-sticks representation. **b,c**, Detailed views of the BIR3 – SERK1 complex interface. Selected interface residues are shown in ball-and-sticks representation with the mutationally analyzed Tyr100, Asp122 and Y214 highlighted in yellow. Water molecules are depicted as red spheres, polar interactions are shown as dotted lines. **d**, ITC binding experiments of BIR2 vs. different SERK3 mutants. **e**, Binding kinetics of SERK3^Y100A^ and SERK3^Y124A^ to BL-associated BRI1 derived from GCI experiments. Fitted kinetic parameters are shown alongside. **f**, Co-immunoprecipitation (Co-IP) experiment using different SERK3 lines vs. BIR3. Input western-blots and a Ponceau stained membrane are shown alongside. **g**, Size-exclusion chromatography experiments using the BIR2, SERK3, BRI1 ectodomains. BIR2 forms no complex with BRI1 (red line), and is not able to dissociate a preformed BRI1 – BL – SERK3 complex (gray line). However, incubation of a preformed BIR2 – SERK3 complex with BRI1 – BL reveals formation of BRI1 – BL – SERK3 complexes (black line), suggesting that BRI1 – BL can compete with BIR2 for SERK3 binding. Void (v_0_) volume and total volume (v_t_) are shown, together with elution volumes for molecular mass standards (Al, Aldolase, 158,000 Da; Ov, Ovalbumin, 44,000 Da; CA, Carbonic anhydrase, 29,000 Da). Peak fractions were analyzed by SDS-PAGE.

An additional set of hydrophobic contacts involving BIR3^W67^ (corresponds to BIR2^W73^ analyzed in Fig. 3c,d), BIR3^I75^, BIR3^Y122^, BIR3^V124^ and BIR3^F146^ (corresponds to BIR2^F152^, see Fig. 3c,d) and SERK residues Val168, Ile192, Pro191 are dominating the interactions between the BIR3 and SERK1 C-terminal halves (Fig. 4a,c). BIR3^R170^, the corresponding mutation in BIR2^R176^ to alanine disrupts complex formation with SERK3 (Fig. 3d), forms hydrogen bonds with backbone atoms in the SERK1 C-cap and other polar contacts are mediated by water molecules (Fig. 4c). Taken together, BIR3 targets the central LRR surface of SERKs normally used for the interaction with ligand-sensing LRR-RKs. The unique protruding loop in BIRs directly contacts the *elg* surface patch, rationalizing the reduced binding of SERK3^D122N^ to BIR ectodomains *in vitro* (Fig. 2a,b).

We next tested if the SERK - BIR LRR domain complex interface controls association of the full-length proteins *in planta.* We found that wild-type SERK3 associated with BIR3 in co-immunoprecipitation experiments (Fig. 4f), as shown previously^17^. The SERK3^D122N^, SERK3^D122A^, SERK3^Y100A^, SERK3^Y124A^ mutants, all of which show reduced binding to isolated BIR LRR domains *in vitro*, consistently show reduced interaction with BIR3 *in vivo* (Fig. 4f). SERK3^F60^ lies outside the SERK - BIR complex interface, but forms part of the BRI1 - SERK steroid binding pocket^5,6^ and its mutation to alanine disrupts BR complex formation *in vitro* and *in planta*^10^. Consistent with our BIR targeting model, the SERK3^F60A^ mutant shows wild-type binding to BIRs in ITC assays and retains interaction with BIR3 *in vivo* (Fig. 4d,f).

Our biochemical observation that SERKs can form tight heterodimeric complexes with BRI1 or with BIRs using largely overlapping interaction surfaces (Fig. S7), prompted us to investigate if the BRI1 and BIR ectodomains could compete for SERK binding. We performed analytical size-exclusion chromatography experiments with the isolated BRI1, SERK3 and BIR2 LRR domains and in the pre- or ab-sence of the steroid hormone. In our ITC assay (Fig. 2a), we could not detect complex formation between BRI1 and BIR3, and consistently BIR2 was unable to dissociate an already formed BRI1-BL-SERK3 complex (Fig. 4g). However, BIR2 could efficiently compete with BRI1 for SERK3 binding (Fig. 4g), in line with our observation that the experimentally determined stoichiometries, binding affinities and -kinetics for the different complexes are similar (Figs. 1d, 2a).

Taken together, the molecular characterization of the SERK3 *elg* allele has revealed that the BR signaling pathway is under negative regulation by the ectodomain of BIR3. We show that SERK3^D122N^ disrupts BIR but not BRI1 binding and thus exhibits a gain-of-function phenotype (Figs. 1c,2b). Mutation of the neighboring SERK3^Y100^ and SERK3^Y124^ to alanine strongly decreases BIR binding, but only SERK3^Y124A^ retains the ability to bind BRI1 - BL with high affinity (Figs. 4d-f). Consistently, SERK3^Y124A^, but not SERK3^Y100A^ or SERK3^Y100A/Y124A^ displays a statistically significant gain-of-function phenotype in hypocotyl growth assays (Fig. 1b,c). The BR-specific nature of the *elg* allele may thus be related to its ability to bind BRI1, but not other SERK3-dependent LRR-RKs with high affinity^12^. The *elg* and *bir3* phenotypes and our quantitative biochemical assays reveal that BRI1 and BIRs can compete for binding to SERKs, with BRI1 being able to out-compete BIRs in the presence of BL. We speculate that this negative regulation of SERKs by BIR proteins may allow for sharper signal transitions, with signaling competent BR complexes forming only in response to significant changes in BR concentration.

Specific physiological functions have been genetically assigned to the different BIR family members in Arabidopsis: BIR1, a catalytically active protein kinase, specifically inhibits SERK3 co-receptor function in immunity and cell death, with *bir1* loss-of-function mutants showing constitutive defense responses associated with a severe growth phenotype^20-22^. BIR2 and BIR3 are additional SERK3 interactors and both proteins are pseudokinases^17,18,23^. Different *bir2* knock-down lines show altered immune responses but no BR signaling phenotype, while *bir3* loss- and gain-of function mutants affect BR signaling (Fig. 2c)^17,18^. We cannot rationalize these specific functions of the different BIRs at the biochemical level, as all BIR ectodomains tested bind various SERK proteins with similar dissociation constants (Fig. 2a), in agreement with a recent study on the role of BIR1 in FLS2-mediated immune signaling^22^. This behavior of BIR proteins is reminiscent of SERKs, which also are largely promiscuous at the biochemical level, but which show partly specific, partly overlapping functions in plant growth, development and immunity^9^. While BIR ectodomains and not their cytosolic kinase domains allow for high affinity SERK binding (Figs. 2-4, S2), BIR signaling specificity may be encoded in their cytosolic domains, as seen with ligand-sensing LRR-RKs^10,24^. In line with this, specific BIR adapter proteins have been reported^25,26^, which could allow for the targeting of BIR family members to specific membrane (nano)-domains^27^, and which could help to create specific signaling outputs in the cytosol^25^. The fact that the *bir3-2* mutant does not phenocopy *elg* plants (Figs. 1b, 2c), suggests that other negative regulators of BR signaling complexes remain to be discovered in the future.

## Methods

### Plant material and growth conditions

Genomic *SERK3* was amplified from *Arabidopsis thaliana* (ecotype Col-0), cloned into pDONR221 (ThermoFisher Scientific) and mutations were introduced by site directed mutagenesis (TableS3). Constructs were assembled employing multi-site Gateway technology into the binary vector pH7m34GW (ThermoFisher Scientific), introduced in the *Agrobacterium tumefaciens* strain pGV2260, and transformed into *Arabidopsis using* the floral dip method^28^. Plants were grown in long day conditions (16 h light) at 21 °C, 50 % humidity and analyzed in homozygous T3 generation. The *bir2-1, bir2-3* and *bir3-2* T-DNA insertion lines were obtained from the Nottingham Arabidopsis Stock Center (NASC).

### Hypocotyl growth assay

After surface sterilization with 70 % ethanol, 0.1 % Triton X-100 for 20 min and stratification at 4 °C for 2 days, seeds were plated on V MS, 0.8 % agar plates supplemented with either 1 μM brassinazole (BRZ, from a 10 mM stock solution in 100 % DMSO, Tokyo Chemical Industry Co. LTD) or, for the controls, with 0.1 % (v/v) DMSO. After light exposure for 1 h, plates were incubated at 22 °C for 5 d in the dark and subsequently scanned at 600 dpi on a regular flatbed scanner (CanoScan 9000F, Canon). Measurements were taken using FIJI^29^ and analyzed with the packages mratios^30^ and multcomp^31^ as implemented in R^32^ (version 3.3.2). We report unadjusted 95% confidence limits for fold-changes instead of p-values^33^. Log-transformed endpoint hypocotyl lengths were analyzed employing a mixed effects model for the ratio of of a given line 'to the wildtype Col-0 allowing heterogeneous variances. To evaluate the treatment-by-mutant interaction, the 95 % two-sided confidence intervals for the relative inhibition (Col-0: untreated vs. BRZ-treated hypocotyl length)/(any genotype: untreated vs. BRZ-treated hypocotyl length) was calculated for the log-transformed length. Hypocotyl growth assays were performed three times, with similar results.

### Protein expression and purification

SERK2^1-220^, SERK3^1-220^ and BRI1^1-799^ were amplified from *A. thaliana* cDNA and BIR1^1-219^, BIR2^1-222^, BIR3^1-213^ and BIR4^1-206^ from *A thaliana* genomic DNA. BIR2^1-222^ was in addition obtained codon-optimized for expression in *Trichoplusia ni* (Tnao), (Invitrogen GeneArt, Germany), SERK1^24-213^ was obtained codon optimized and fused to an azurocidin signal peptide; all constructs were cloned in a modified pFastBac vector (Geneva Biotech), containing a TEV (tobacco etch virus protease) cleavable C-terminal StrepII-9xHis tag. Mutations were created using site directed mutagenesis (TableS3). Tnao38^34^ cells were infected with a multiplicity of infection (MOI) of 1 for SERKs or 3 for BRI1 and BIRs at a density of 2x10^6^cells/ml and incubated 26 h at 28 °C and 48 h at 22 °C. Subsequently the secreted proteins were purified from the supernatant by Ni^2+^ (HisTrap excel; GE Healthcare; equilibrated in 25 mM KP_i_ pH 7.8, 500 mM NaCl) and StrepII (Strep-Tactin Superflow high capacity; IBA; equilibrated in 25 mM Tris pH 8.0, 250 mM NaCl, 1 mM EDTA) affinity chromatography. The purity of the preparations was further improved by size-exclusion chromatography on either a Superdex 200 increase 10/300 GL, HiLoad 16/600 Superdex 200 pg or HiLoad 26/600 Superdex 200 pg column (GE Healthcare), equilibrated in 20 mM sodium citrate pH 5.0, 150 mM NaCl.

The cytosolic domain of BIR2 (residues 258-605 or 289-605) was cloned in a modified pET vector (Novagen) providing a TEV cleavable N-terminal 8xHis-StrepII-Thioredoxin tag, constructs were transformed in *E.coli* BL21 (DE3) RIL cells. Protein expression was induced by adding IPTG (0.5 mM final concentration) to cell cultures grown at 37 °C to a OD_600_= 0.6 and bacteria were harvested after incubation for 18 h at 16 °C. SERK3 (residues 250-615) and BRI1 (residues 814-1196) cytoplasmic domains were cloned in a modified pFastBac vector (Geneva Biotech) with a TEV-cleavable N-terminal 10xHis-2xStrepII tag for expression in insect cells. Proteins were expressed in Tnao38 cells for three days at 28 °C after infection with a MOI of 2.

For purification from bacterial as well as from insect cells, pellets were resuspended in buffer A (20 mM Hepes pH 7.5, 500 mM NaCl, 4 mM MgCl2, 2mM β-mercaptoethanol) and disrupted by sonication. The cell debris was removed by centrifugation at 20,000 g for 1 h at 4 °C and the recombinant proteins were purified by sequential Ni^2+^ (HisTrap excel; GE Healthcare; equilibrated in buffer A) and StrepII (Strep-Tactin XT Superflow; IBA; equilibrated in 25 mM Tris pH 8.0, 250 mM NaCl, 1 mM EDTA) affinity chromatography. The tags were cleaved-off by incubating the protein with TEV protease overnight at 4 °C. The cleaved tags and the protease were removed by an additional Ni^2+^ affinity chromatography step. The recombinant proteins were further purified by size exclusion chromatography at 4 °C on a HiLoad 16/600 Superdex 200 pg (GE Healthcare) equilibrated with 20 mM Hepes pH 7.5, 150 mM NaCl, 1 mM MgCh, 0.5 mM TCEP. Proteins concentrated to 15 mg/ml and snap frozen in liquid N2.

Molar protein concentrations for BIR2, BIR3, SERK1, SERK3 and BRI1 were calculated using their molar extinction coefficient and molecular weights of 23.4, 24.0, 25.2, 27.4, 105.0 kDa, respectively (as determined by MALDI-TOF mass spectrometry). The calculated molecular masses for BIR2^289-605^, BIR2^258-605^, SERK3^250-615^, BRI1^814^'^1196^ are 35.3, 38.8, 41.5, 42.7 kDa, respectively.

### Grating coupled interferometry (GCI)

The Creoptix WAVE system (Creoptix AG, Switzerland), a label-free surface biosensor^35^ was used to perform GCI experiments. All experiments were performed on 2PCP WAVEchips (quasi-planar polycarboxylate surface; Creoptix AG, Switzerland). After a borate buffer conditioning (100 mM sodium borate pH 9.0, 1 M NaCl; Xantec, Germany) the respective LRR ectodomain was immobilized on the chip surface using standard amine-coupling: 7 min activation (1:1 mix of 400 mM N-(3-dimethylaminopropyl)-N’-ethylcarbodiimide hydrochloride and 100 mM N-hydroxysuccinimide [both Xantec, Germany]), injection of the LRR domain (10 to 40 μg/ml) in 10 mM sodium acetate pH 5.0 (Sigma, Germany) until the desired density was reached, passivation of the surface (0.5% BSA [Roche, Switzerland] in 10mM sodium acetate pH 5.0) and final quenching with 1 M ethanolamine pH 8.0 for 7 min (Xantec, Germany). For a typical experiment, SERK3 was injected in a 1:2 dilution series (starting from 2 μM) in 20mM citrate pH 5.0, 250mM NaCl at 25°C.

Blank injections were used for double referencing and a DMSO calibration curve for bulk correction. Analysis and correction of the obtained data was performed using the Creoptix WAVEcontrol software (applied corrections: X and Y offset, DMSO calibration, double referencing) and a one-to-one binding model with bulk correction was used to fit all experiments.

### Isothermal titration calorimetry (ITC)

All ITC experiments were performed on a Nano ITC (TA Instruments) with a 1.0 ml standard cell and a 250 μl titration syringe at 25 °C. Proteins were gelfiltrated or dialyzed into ITC buffer (20 mM sodium citrate pH 5.0, 150 mM NaCl for LRR domains / 20 mM Hepes pH 7.5, 150 mM NaCl, 1 mM MgCl_2_, 0.5 mM TCEP for kinase domains) prior to all experiments. For a typical ectodomain experiment, 16 μl of BIR (at ~400 μM) was injected into ~40 μM SERK protein in the cell at 150 s intervals (15 injections). Experiments with the kinase domains were performed by injecting 10 μl of BIR2 or BRI1 cytosolic domain at ~200 μM into ~20 μM of SERK3 kinase domain in the cell at 150s intervals (25 injections). Data was corrected for the dilution heat and analyzed using NanoAnalyze program (version 3.5) as provided by the manufacturer. All quantitative biochemical assays were performed at least twice.

### Protein crystallization and data collection

Crystals of the isolated BIR2 ectodomain were grown in sitting drops composed of 0.2 μl of protein solution (BIR2^1-222^ at 9 mg/ml in 20 mM sodium citrate pH 5.0, 150 mM NaCl) and 0.2 μl of 1.8 M sodium malonate pH 4.0. Crystals formed after several months, were cryoprotected in 2.4 M sodium malonate pH 4.0 and were snap frozen in liquid N_2_. Native (λ= 1.000020 Å) and anomalous (λ= 1.999770 Å) datasets were collected from a single crystal at beam line PX-III of the Swiss Light Source, Villigen (Table S2). Crystals of the BIR3^1-213^ - SERK1^24-213^ complex were grown from hanging drops containing 1 μl of protein solution (14 mg/ml in 20 mM sodium citrate pH 5.0, 150 mM NaCl) and crystallization buffer (19% [v/v] PEG 3,350, 1M LiCl, 0.1 M sodium acetate pH 5.5), suspended over 0.6 ml of the latter as reservoir solution. Crystals were cryoprotected by serial transfer in reservoir solution supplemented with a final concentration of 15% (v/v) glycerol. Crystals diffracted up to 1.0 Å at PX-III and due to the beam line geometry, a complete dataset at 1.25 Å was recorded (Table S2). Data processing and scaling was done with XDS^36^ (version: June, 2017).

### Crystallographic structure solution and refinement

The BIR2 anomalous dataset was used for experimental phasing using the Single Anomalous Diffraction (SAD) method. Ten consistent sulfur sites were identified using ShelxD^37^ and Phenix.hyss^38^ and used for site refinement and phasing in Sharp^39^ (Table S2). Density modification, 2-fold NCS averaging and phase extension to 1.9 Å in the program Phenix.resolve^40^ yielded a readily interpretable electron density map and the structure was completed in alternating cycles of manual building/rebuilding in Coot^41^, and restrained TLS refinement in Refmac5^42^.

The structure of the BIR3 - SERK1 complex was solved using the molecular replacement method as implemented in the program Phaser^43^, and using the isolated BIR2 and SERK1 (PDB-ID 4LSC^5^) structures as search models. The solution comprises a hetero-dimer in the asymmetric unit and the structure was completed by manual correction in Coot and anisotropic refinement in Refmac5.

The quality of the refined structures were assessed using the program Molprobity^44^, structural diagrams were made with Pymol (https://sourceforge.net/projects/pymol/) and Chimera^45^.

### Analytical size exclusion chromatography

Gel filtration experiments were performed using a Superdex 200 Increase 10/300 GL column (GE Healthcare) pre-equilibrated in either 20 mM sodium citrate pH 5.0, 150 mM NaCl for LRR domain interaction assays, or with 20 mM Hepes pH 7.5, 150 mM NaCl, 1 mM MgCl_2_, 0.5 mM TCEP for cytplasmic domain oligomeric state analysis. 500 μl of the respective protein (0.2 mg/mL) was loaded sequentially onto the column and elution at 0.75 ml/min was monitored by ultraviolet absorbance at 280 nm. BL concentration was 1 μM in the BRI1 - BL - SERK3 complex sample prior to loading.

### Plant protein extraction and immunoprecipitation

Surface-sterilized and stratified seeds were plated on V MS, 0.8 % agar plates and grown for ~14 d. Seedlings were frozen in liquid N2, ground to fine powder using mortar and pestel (1 g per sample) and resuspended in 3 ml of ice cold extraction buffer (50 mM Bis-Tris pH 7.0, 150mM NaCl, 10 % (v/v) glycerol, 1 % Triton X-100, 5 mM DTT, protease inhibitor cocktail (P9599, Sigma). After gentle agitation for 1 h at 4 °C, samples were centrifuged for 30 min at 4 °C and 16,000 g; the supernatant was transferred to a fresh tube and the protein concentration measured using a Bradford assay. 20 mg of total protein in a volume of 5 ml were incubated with 50 μl of anti-HA superparamagnetic MicroBeads (Miltenyi Biotec) for 1 h at 4 °C with agitation for each co-immunoprecipitation (Co-IP). The beads were then collected using μMACS Columns (Miltenyi Biotec), washed 4 times with 1 ml of cold extraction buffer and proteins were eluted in 20+20 μl of extraction buffer at 95 °C. Samples were separated on 10 % SDS-PAGE gels; In the subsequent western blots SERK3:6HA was detected using anti-HA antibody coupled to horse radish peroxidase (HRP, Miltenyi Biotec) at 1:5,000 dilution, while BIR3 was detected using a polyclonal BIR3 antibody^17^ at 1:500 dilution followed a secondary anti-rabbit HRP antibody (1:10,000, Calbiochem #401353). Co-immunoprecipitation experiments were repeated two times, with similar outcome.

## Acknowledgements

We thank B. Kemmerling for kindly providing us with BIR2 and BIR3 polyclonal antibodies, staff at beam line PXIII of the Swiss Light Source, Villigen, Switzerland for technical assistance during data collection, J. Santiago for providing the SERK2 expression plasmid, and K. Lau for help with preparing figures. Crystallographic coordinates and structure factors have been deposited with the Protein Data Bank (http://rcsb.org) with accession codes 6FG7 (BIR2) and 6FG8 (BIR3 - SERK1). This work was supported by grant 31003A_176237 from the Swiss National Science Foundation and by an International Research Scholar Award from the Howard Hughes Medical Institute (to MH).

## Supplemental Figures

**Fig. S1:**
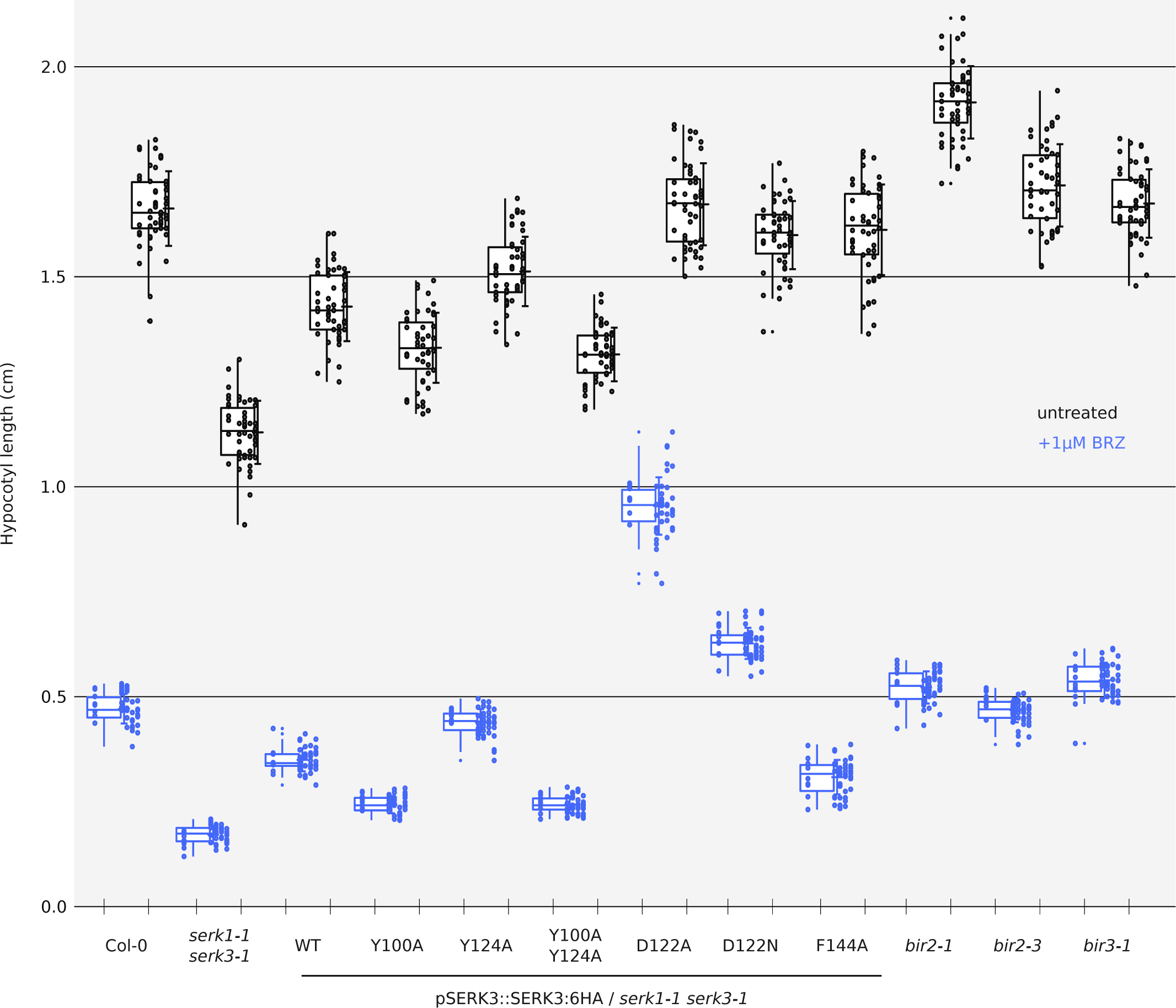
Hypocotyl growth assay raw data. Shown are box plots with the raw data depicted as individual dots. Untreated: black, BRZ treated: blue, N=5, n=50.

**Fig. S2:**
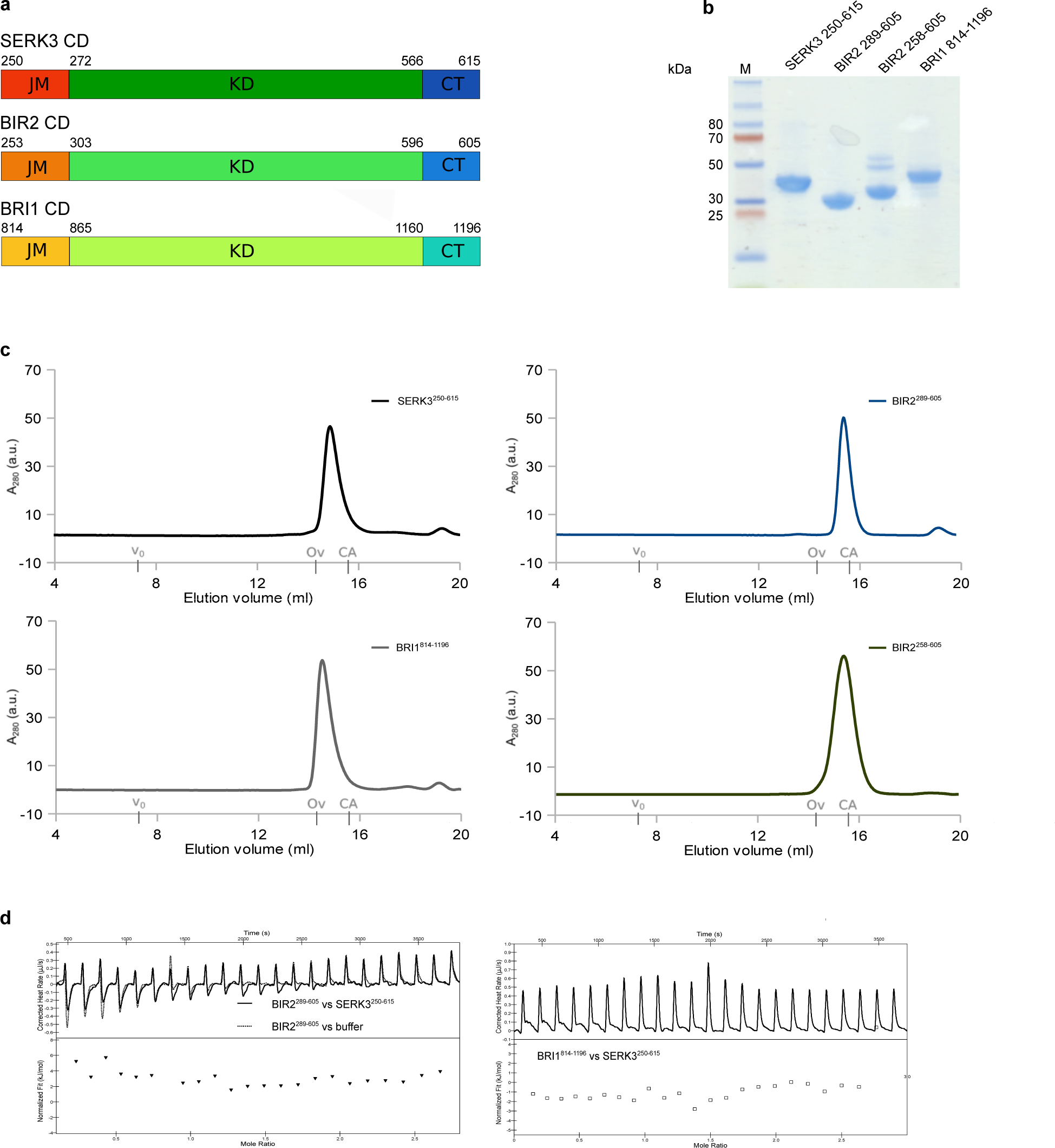
The recombinant BIR2 and SERK3 cytoplasmic domains do not interact *in vitro*. **a**, Structural organization of the SERK3, BIR2 and BRI1 cytoplasmic domains (CD) with domain borders included. JM, juxtamembrane domain; KD, kinase domain; CT, C-terminal domain. **b,c**, Analysis of the purified cytoplasmic domains on (**b**) a Coomassie stained 10 % SDS-PAGE gel and (**c**) by size exclusion chromatography on a Superdex 200 increase 10/300 GL column (GE Healthcare) reveals that all isolated cytoplasmic domains behave as apparent monomers in solution. The void (v_0_) volume is shown, together with elution volumes for molecular mass standards (Ov, Ovalbumin, 44,000 Da; CA, Carbonic anhydrase, 29,000 Da). **d**, Isothermal titration calorimetry (ITC) experiments with cytoplasmic domains of SERK3 vs. BIR2 (left) and BRI1 (right). No binding was detected, suggesting that the binding affinity between BIR2 and SERK3 or BRI1 and SERK3 is relatively low. Thus, BIR binding may be driven by their extracellular, rather than by their cytoplasmic domains.

**Fig. S3:**
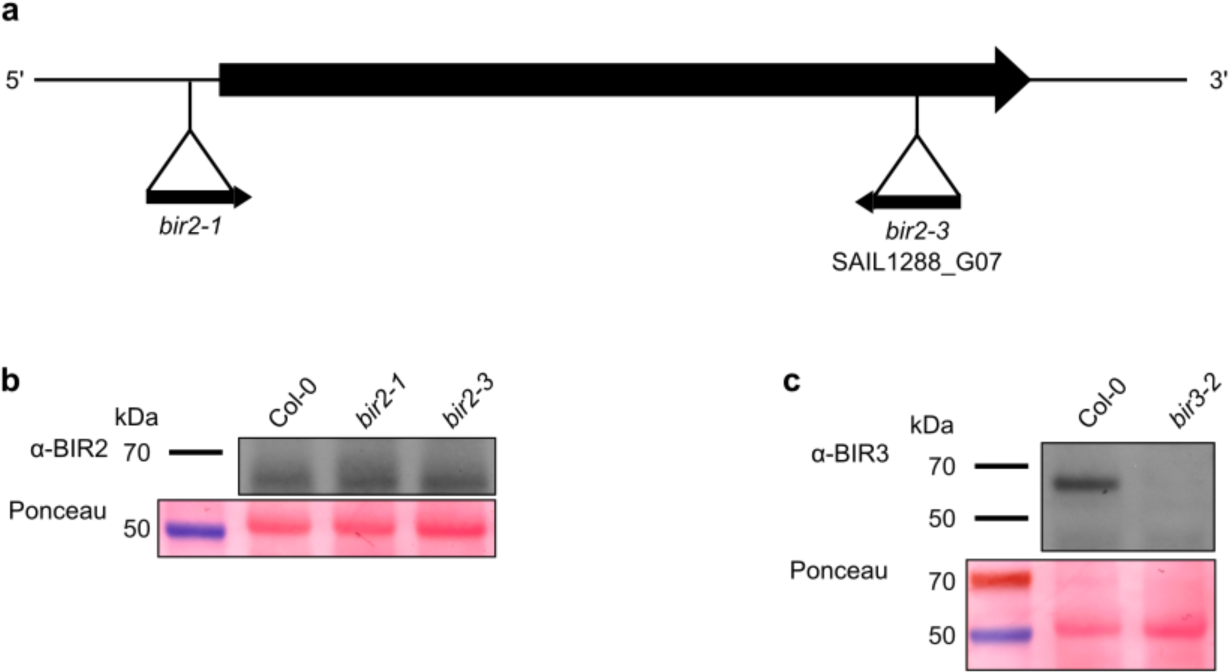
Expression levels of BIR2/3 in mutant lines. **a**, Schematic overview of the T-DNA insertion sites *bir2-1*^18^ and *bir2-3* (SAIL1288_G07, this study) shown as black triangles in the *BIR2* locus (bold black arrow). **b**, Analysis of BIR2 protein levels in wild-type Col-0, *bir2-1* and *bir2-3* mutant plants. **c**, BIR3 protein levels in Col-0 and *bir3-2*^17^ mutant lines. Ponceau stained membranes are shown alongside as loading controls.

**Fig. S4:**
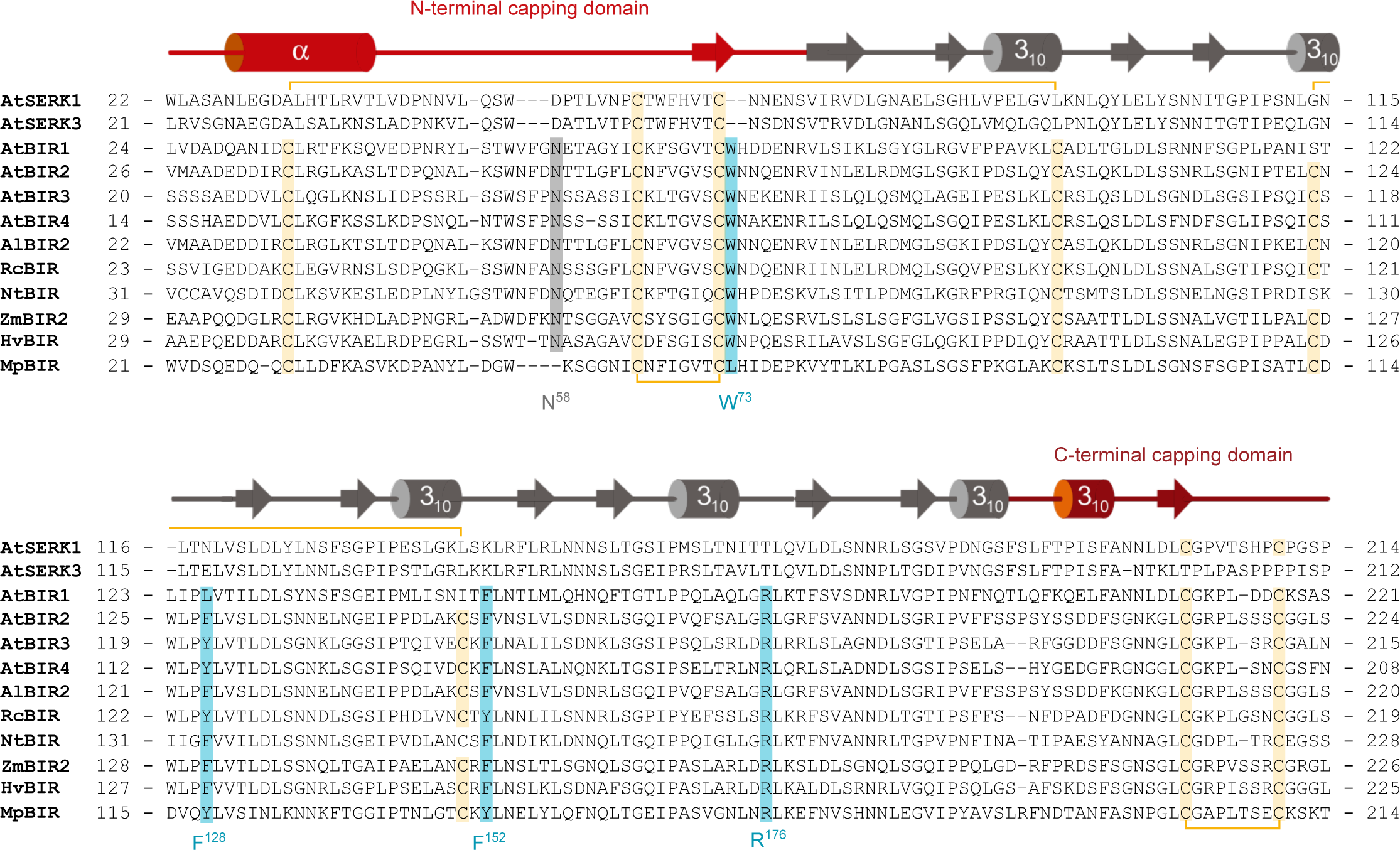
BIR – SERK complex interface residues are conserved among BIR family members from different species. **a**, Structure based sequence alignment of the ectodomains of *Arabidopsis thaliana* SERK1 (Uniprot [http://www.uniprot.org] identifier: Q94AG2), SERK3 (Uniprot identifier: Q94F62), BIR1 (Uniprot identifier: Q9ASS4), BIR2 (Uniprot identifier: Q9LSI9), BIR3 (Uniprot identifier: O04567), BIR4 (Uniprot identifier: C0LGI5), *Arabidopsis lyrata* BIR2 (Uniprot identifier: D7LPU1), *Ricinus communis* BIR (Uniprot identifier: B9RUI5), *Nicotiana tabacum* BIR (Uniprot identifier: A0A1S4BB12), *Zea mays* BIR2 (Uniprot identifier: K7TUC5), *Hordeum vulgare* BIR (Uniprot identifier: F2E7N3) and *Marchantia polymorpha* BIR (Uniprot identifier: A7VM20). Shown alongside is a secondary structure assignment, with the N- and C-terminal capping domains highlighted in red, calculated using DSSP^46^. BIR residues of the lateral protein interaction interface are highlighted in blue, disulfide bridges in yellow and the conserved N-terminal glycosylation site in gray. All numbering refers to AtBIR2.

**Fig. S5:**
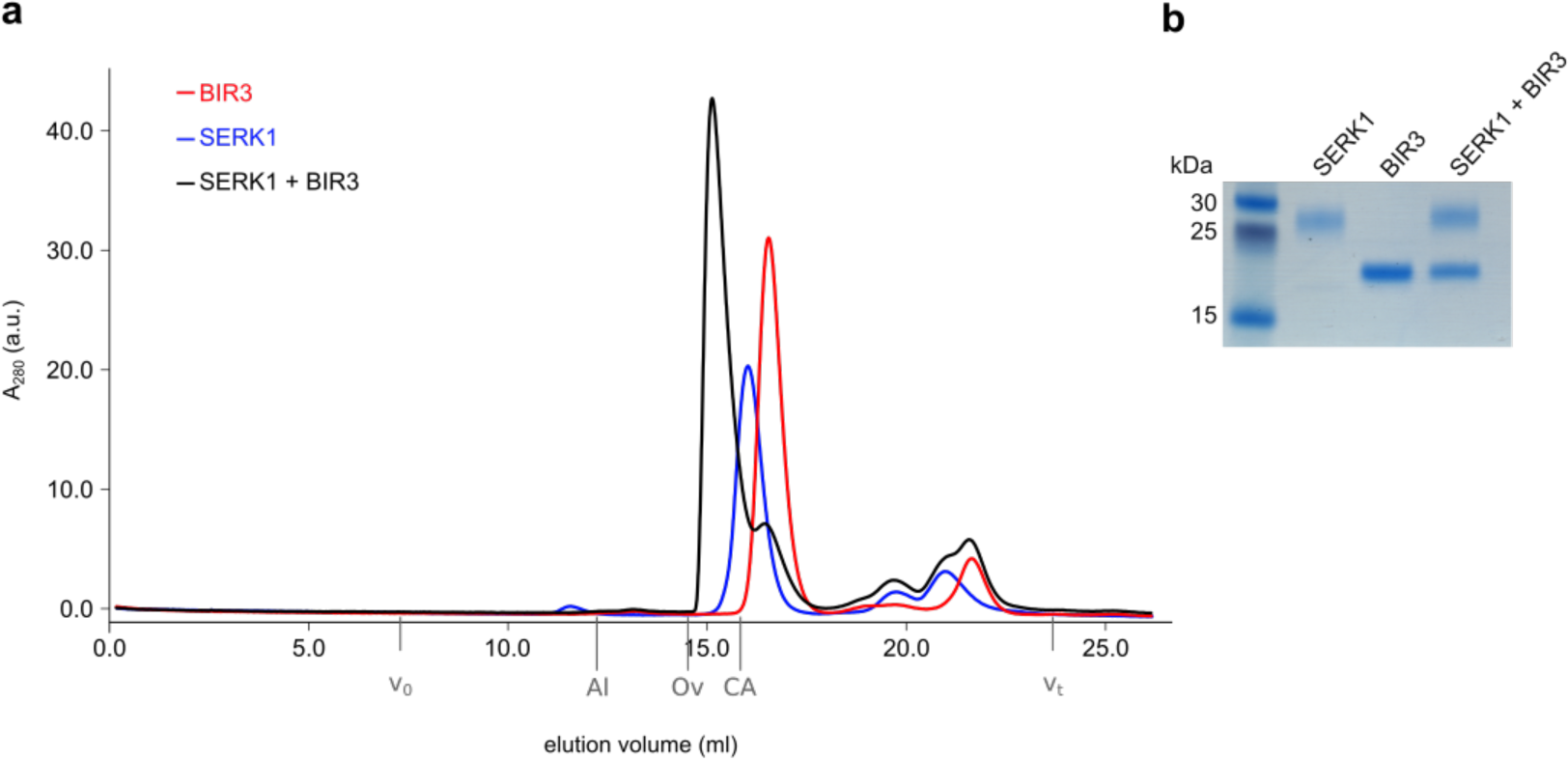
The BIR3 and SERK1 ectodomains form heterodimers in solution. **a,b**, Analytical size exclusion chromatography. The isolated BIR3 (red absorption trace) and SERK3 (blue) ectodomains elute as apparent monomers when run in isolation, and form a heterodimeric complex (black line). Void (v_0_) volume and total volume (v_t_) are shown, together with elution volumes for molecular mass standards (Al, Aldolase, 158,000 Da; Ov, Ovalbumin, 44,000 Da; CA, Carbonic anhydrase, 29,000 Da). A SDS PAGE analysis of the peak fractions is shown in (**b**).

**Fig. S6:**
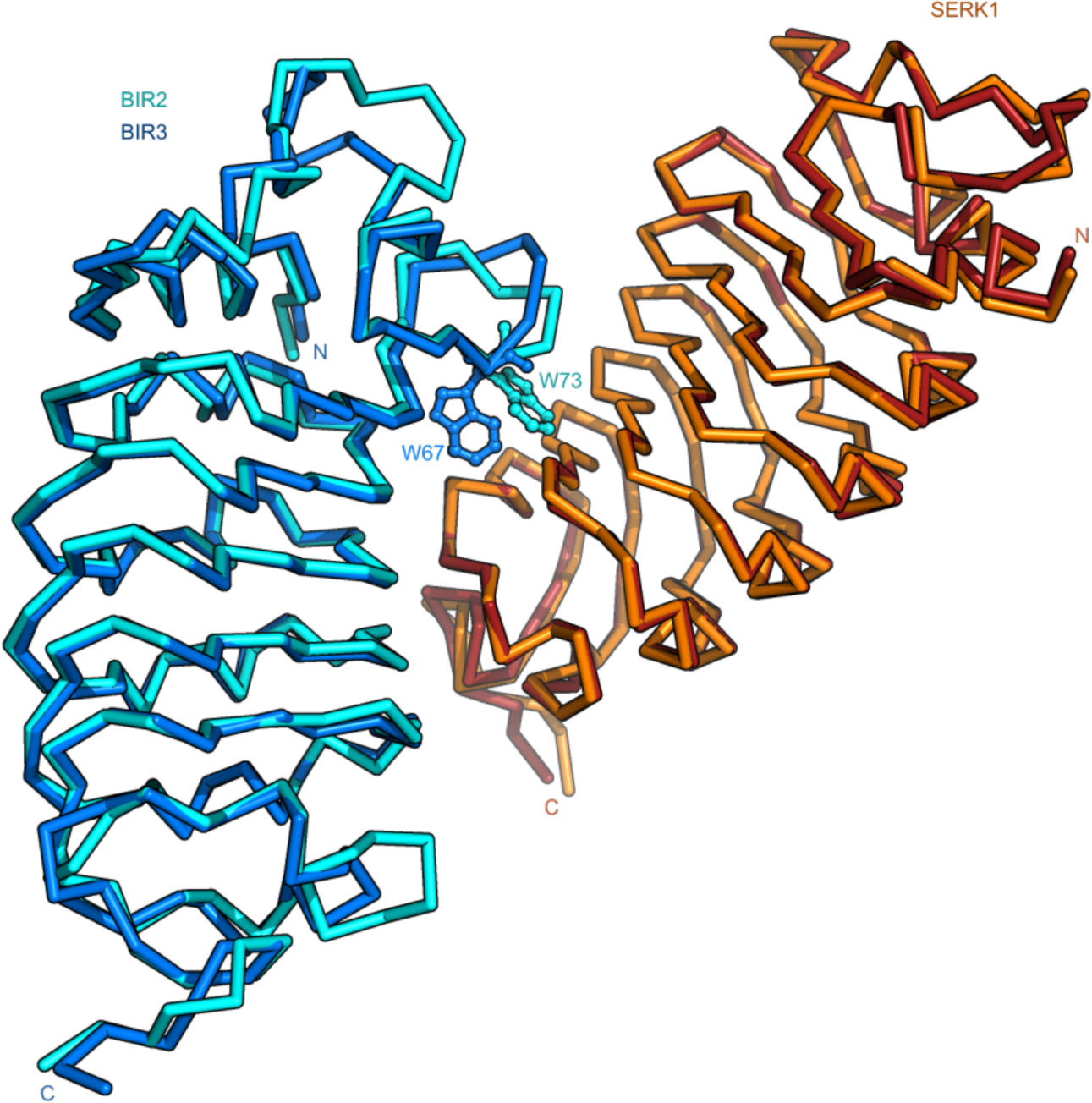
No major conformational changes occur upon BIR3 – SERK1 complex formation. Structural superposition of the BIR3 – SERK1 complex with the isolated BIR2 (r.m.s.d. is ~1.2 Å comparing 160 corresponding C_α_ atoms) and SERK1 (PDB-ID 4LSC^5^, r.m.s.d. is ~0.9 Å comparing 186 corresponding C_α_ atoms) ectodomains. Shown are C_α_ traces of SERK1 (orange for the isolated ectodomain and red for SERK1 in complex with BIR3), BIR2 (in cyan) and BIR3 (in blue). BIR3W67 and the corresponding BIR2W73 are highlighted as ball-and-sticks.

**Fig. S7:**
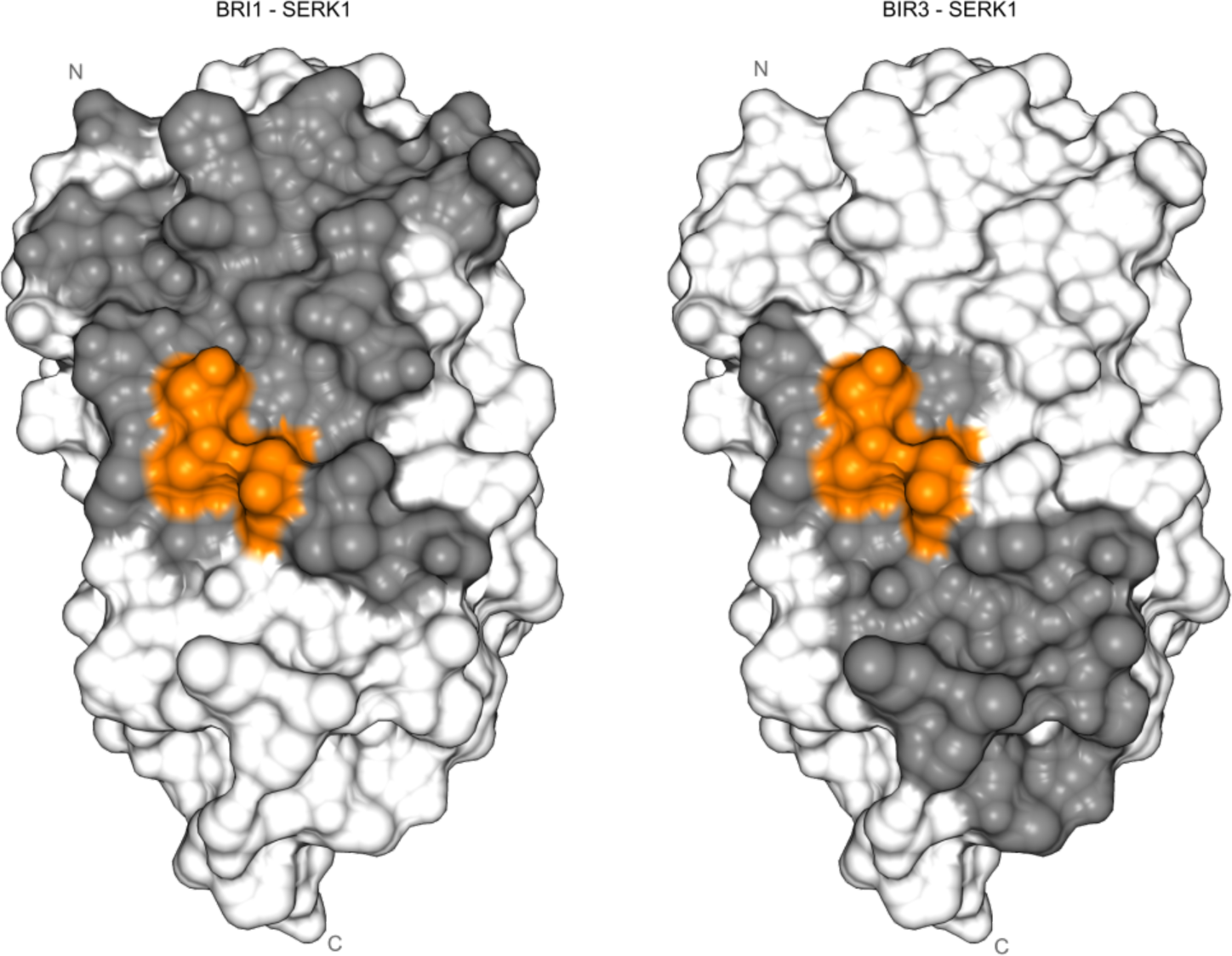
Partly overlapping surface areas in SERK1 are involved in BRI1 and BIR3 binding, respectively. Surface view of the SERK1 ectodomain with BRI1 (left) and BIR3 (right) interacting residues (defined using the program PISA^47^) shown in dark gray. Interaction with BRI1 involves mainly residues originating from the SERK1 N-terminal cap, while the interaction with BIR3 involves residues from the two C-terminal LRRs and from the C-terminal cap. Importantly, the *elg* mutation and the corresponding SERK3^D122^ forms part of both complex interfaces (highlighted in orange).

**Table S1:**
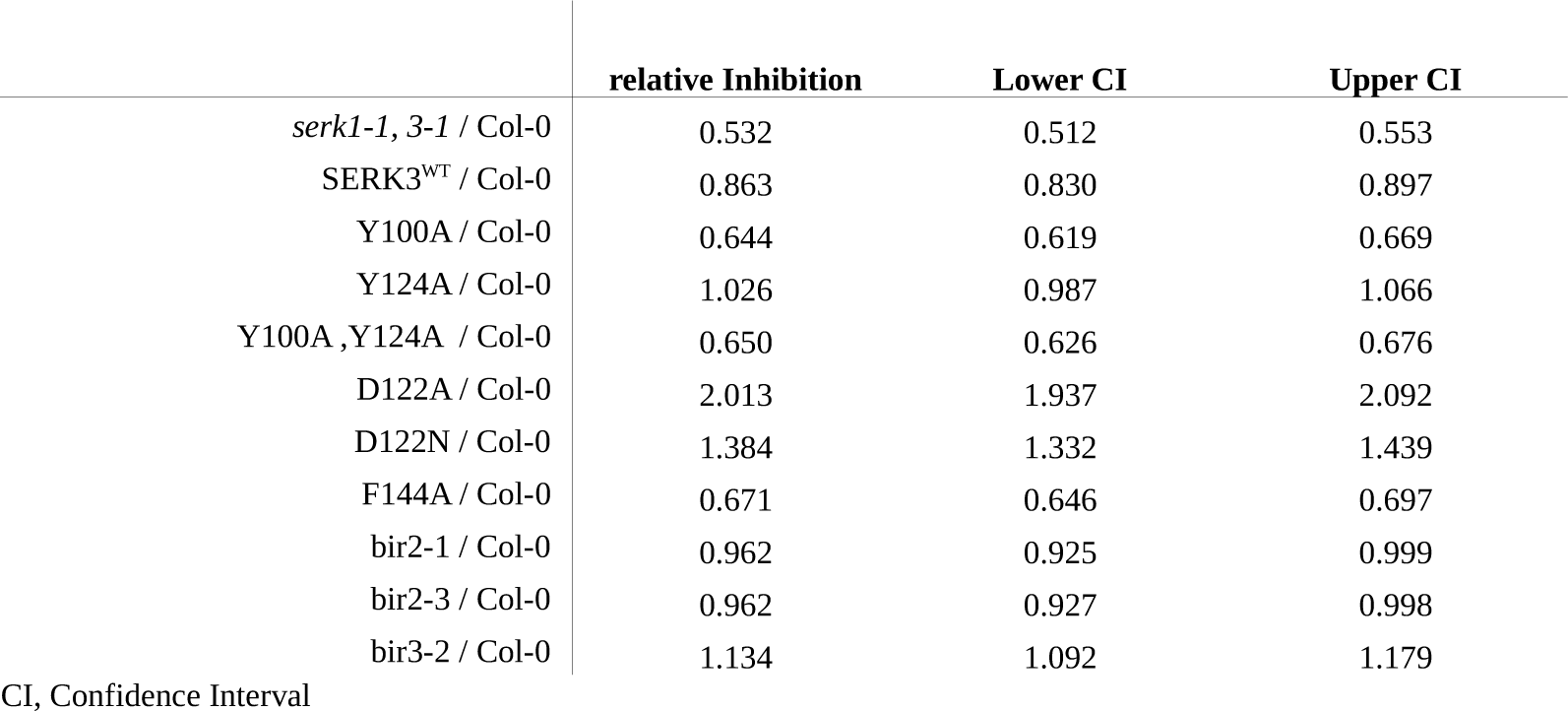
Statistical evaluation of the hypocotyl growth assay

|  | relative Inhibition | Lower CI | Upper CI |
| --- | --- | --- | --- |
| *serk1-1, 3-1* / Col-0 | 0.532 | 0.512 | 0.553 |
| SERK3<sup>WT</sup> / Col-0 | 0.863 | 0.830 | 0.897 |
| Y100A / Col-0 | 0.644 | 0.619 | 0.669 |
| Y124A / Col-0 | 1.026 | 0.987 | 1.066 |
| Y100A,Y124A / Col-0 | 0.650 | 0.626 | 0.676 |
| D122A / Col-0 | 2.013 | 1.937 | 2.092 |
| D122N / Col-0 | 1.384 | 1.332 | 1.439 |
| F144A / Col-0 | 0.671 | 0.646 | 0.697 |
| bir2-1 / Col-0 | 0.962 | 0.925 | 0.999 |
| bir2-3 / Col-0 | 0.962 | 0.927 | 0.998 |
| bir3-2 / Col-0 | 1.134 | 1.092 | 1.179 |

CI, Confidence Interval

**Table S2.** Crystallographic data collection, phasing and refinement.

|  | BIR2<br>*sulfur SAD* | BIR2<br>*native* | BIR3 – SERK1<br>*native* |
| --- | --- | --- | --- |
| **Data collection** |  |  |  |
| Space group | *P*6<sub>4</sub>22 | *P*6<sub>4</sub>22 | *P*2<sub>1</sub> |
| Wavelength (Å) | 1.999770 | 1.000020 | 1.033201 |
| Cell dimensions |  |  |  |
| *a*, *b*, *c* (Å) | 153.77, 153.77, 110.06 | 153.77, 153.77, 110.06 | 52.17, 50.76, 77.43 |
| $\alpha$, $\beta$, $\gamma$ (°) | 90, 90, 120 | 90, 90, 120 | 90, 96.72, 90 |
| Resolution (Å) | 44.75 – 3.0 (3.08 – 3.0) | 45.77 – 1.90 (2.02 – 1.90) | 40.81 – 1.25 (1.33 – 1.25) |
| *R*<sub>meas</sub><sup>#</sup> | 0.229 (1.00) | 0.221 (2.88) | 0.058 (1.10) |
| CC(1/2) (%)<sup>#</sup> | 100.0 (96.1) | 100.0 (42.9) | 100.0 (85.3) |
| *I*/σ*I*<sup>#</sup> | 35.50 (7.69) | 11.90 (1.0) | 15.1 (1.5) |
| Completeness (%)<sup>#</sup> | 100.0 (99.8) | 100.0 (97.9) | 99.8 (95.5) |
| Redundancy<sup>#</sup> | 121.9 (121.4) | 13.15 (12.8) | 6.3 (5.7) |
| Wilson B-factor<sup>#</sup> |  | 35.0 | 21.4 |
| **Phasing** |  |  |  |
| Resolution (Å) | 44.75 – 3.00 |  |  |
| No. of sites | 10 |  |  |
| Phasing power (ano)<sup>†</sup> | 0.558 |  |  |
| FOM<sup>†</sup> | 0.27 |  |  |
| **Refinement** |  |  |  |
| Resolution (Å) |  | 45.77 – 1.90 | 40.81 – 1.25 |
| No. reflections |  | 57,323 | 104,302 |
| *R*<sub>work</sub>/*R*<sub>free</sub><sup>§</sup> |  | 0.21/0.23 | 0.15/0.18 |
| No. atoms |  |  |  |
| protein |  | 2,986 | 2,962 |
| glycan |  | 59 | 189 |
| PEG |  |  | 44 |
| solvent |  | 120 | 433 |
| Res. B-factors<sup>§</sup> |  |  |  |
| protein |  | 37.4 | 21.1 |
| glycan |  | 54.6 | 72.32 |
| PEG |  |  | 44.8 |
| solvent |  | 36.7 | 39.22 |
| R.m.s deviations<sup>§</sup> |  |  |  |
| Bond lengths (Å) |  | 0.008 | 0.012 |
| Bond angles (°) |  | 1.34 | 1.61 |
| Molprobrity results |  |  |  |
| Ramachandran outliers (%)<sup>‡</sup> |  | 0.0 | 0.0 |
| Ramachandran favored (%)<sup>‡</sup> |  | 95.5 | 95.7 |
| Molprobrity score<sup>‡</sup> |  | 1.32 | 1.33 |
| PDB - ID |  | **6FG7** | **6FG8** |

<sup>#</sup>as defined XDS<sup>36</sup>, <sup>†</sup>in Sharp<sup>39</sup>, <sup>§</sup>in Refmac<sup>542</sup>, or <sup>‡</sup>in Molprobrity<sup>44</sup>, respectively.

**Table S3:**
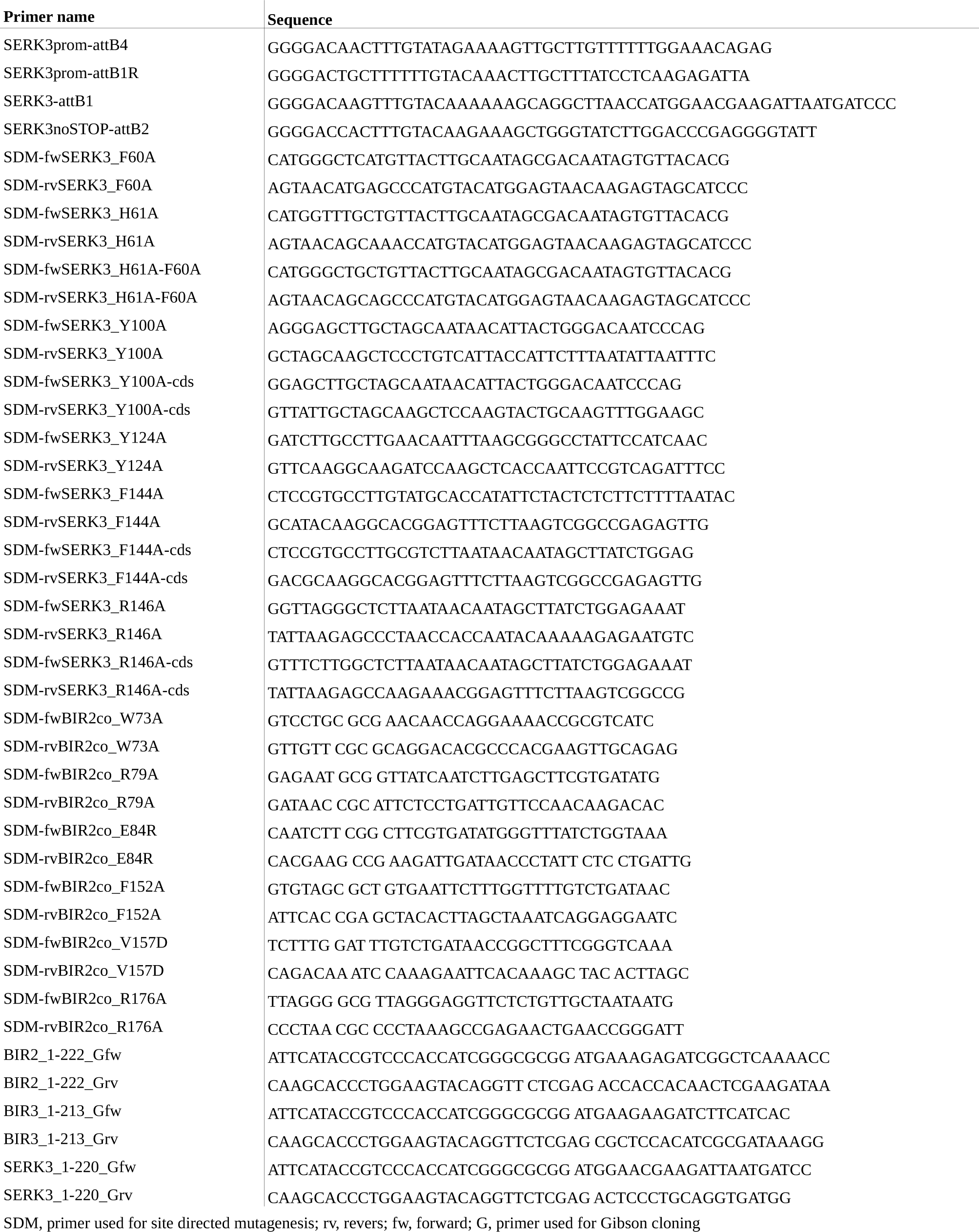
Primers used in this study

